# Generation of nonlinear and spatially-organized 3D cultures on a microfluidic chip using photoreactive thiol-ene and methacryloyl hydrogels

**DOI:** 10.1101/2020.09.09.287870

**Authors:** Jennifer E. Ortiz-Cárdenas, Jonathan M. Zatorski, Abhinav Arneja, Alyssa N. Montalbine, Jennifer M. Munson, Chance John Luckey, Rebecca R. Pompano

**Affiliations:** Department of Chemistry, University of Virginia, PO BOX 400319, Charlottesville, VA, USA 22904; Department of Pathology, University of Virginia, Charlottesville, VA, USA 22904; Department of Biomedical Engineering and Mechanics, Fralin Biomedical Research Institute at Virginia Tech-Carilion, Virginia Polytechnic Institute and State University, Roanoke, VA, USA; Department of Chemistry, Carter Immunology Center, University of Virginia, PO BOX 400319, Charlottesville, VA, USA 22904

**Keywords:** GelMA, GelNB, methacrylate, photopolymerization, organs-on-chip, lymphocytes

## Abstract

Micropatterning techniques for 3D cell cultures enable the recreation of tissue-level structures, but their combination with well-defined, microscale fluidic systems for perfusion remains challenging. To address this technological gap, we developed a user-friendly in-situ micropatterning protocol that integrates photolithography of crosslinkable, cell-laden hydrogels with a simple microfluidic housing, and tested the impact of crosslinking chemistry on stability and spatial resolution. Working with gelatin functionalized with photo-crosslinkable moieties, we found that inclusion of cells at high densities (≥ 10^7^/mL) during crosslinking did not impede thiol-norbornene gelation, but decreased the storage moduli of methacryloyl hydrogels. Hydrogel composition and light dose were selected to match the storage moduli of soft tissues. The cell-laden precursor solution was flowed into a microfluidic chamber and exposed to 405 nm light through a photomask to generate the desired pattern. The on-chip 3D cultures were self-standing, and the designs were interchangeable by simply swapping out the photomask. Thiol-ene hydrogels yielded highly accurate feature sizes from 100 – 900 μm in diameter, whereas methacryloyl hydrogels yielded slightly enlarged features. Furthermore, only thiol-ene hydrogels were mechanically stable under perfusion overnight. Repeated patterning readily generated multi-region cultures, either separately or adjacent, including non-linear boundaries that are challenging to obtain on-chip. As a proof-of-principle, primary human T cells, were patterned on-chip with high regional specificity. Viability remained high (> 85%) after overnight culture with constant perfusion. We envision that this technology will enable researchers to pattern 3D cultures under fluidic control in biomimetic geometries that were previously difficult to obtain.

## I. Introduction

Micropatterned cell cultures have become valuable tools to study cell and tissue behavior, enabling both fundamental mechanistic studies and high-throughput screening of arrays of cell cultures.^1,2^ Whereas optimization of 3D cultures traditionally focuses on the mechanical and biochemical properties of the cell culture matrix, the spatial arrangement of cell-laden 3D culture matrices offers an additional point of control to increase the biomimicry and utility of engineered tissues. For example, micropatterned 3D cultures have been used to replicate emergent tissue-level properties such as tissue hypoxia,^3,4^ cell migratory and homing activity,^5,6^ formation of signaling gradients,^7,8^ and to form cell arrays for high-throughput assays.^9–11^ Integration of spatially organized 3D cultures with platforms that enable fluidic control, such as microfluidic chips, is often advantageous, e.g. for control over media perfusion, but remains challenging.

Standard technologies are limited in their ability to create complex spatial structures under fluidic control, and thus cannot fully replicate the complex spatial organization of native tissues or allow full freedom of high-throughput array design. Integration of 3D cultures with microfluidics is often accomplished by direct micropatterning inside a microfluidic chip, usually through the use of laminar flow and/or physical support structures. Patterns achieved through laminar flow are highly linear, producing well-controlled lanes of hydrogel.^12–15^ Micropillars and other physical supports allow more flexibility by patterning via surface tension, but typically also produce linear or gently curved boundaries.^16^ Both patterning strategies struggle to generate free-standing islands, concentric features, or two or more closely abutting cultures. Furthermore, these patterning strategies largely rely on pre-determined chip geometries, so major changes to the pattern require time-consuming fabrication of new masters.

A potential strategy to overcome these limitations is photolithography, in which light passes through a photomask to pattern a photo-crosslinkable culture matrix.^17^ A significant benefit of photopatterning is the ability to modulate the mechanical properties of the hydrogel, e.g. to match those of a particular tissue, by optimizing the chemical composition of the gel and dose of light.^18–22^ Photolithography is most commonly used to pattern hydrogels onto coverslips or other substrates that are rarely integrated with a flow control system.^23–26^ More rarely, in-situ photolithography has been used to coat the interior of microfluidic channels,^13^ create monolayers of hydrogel onto which cells are later seeded,^27^ and to create free-floating microstructures for collection downstream.^28,29^ Recently, on-chip photolithography was used to create cell-laden micropillar arrays and organized tumor-on-chip cultures.^6,9,30^ However, the impact of materials choice on reproducibility and mechanical stability of cell-laden hydrogels on-chip, remains unexplored.

Patterning cultures on-chip requires crosslinking cell-laden hydrogels without cytotoxicity, while still achieving biomimetic mechanical properties and stability under fluid flow. These requirements impose constraints on the choice of biomaterial and extent of photoexposure. Indeed, the risk of phototoxicity has largely limited the use of photopatterning to hydrogels without cells or with hardy cell lines.^6,22,30–34^ In the presence of a suitable photoinitiator and exposure to UV or violet-blue light, methacrylate- and methacrylamide-functionalized biomaterials undergo chain growth polymerization.^19^ This methacryloyl-based polymerization is common, but is severely inhibited by oxygen and can generate high concentrations of reactive oxygen species. Thus, in the oxygen-rich culture matrix and oxygen-permeable microfluidic devices, such crosslinking chemistry may be toxic. However, newer crosslinking chemistries and photoinitiators have proven less cytotoxic, particularly thiol-norbornene (thiol-ene) chemistry.^21,35^ When crosslinked, thiol- and norbornene-functionalized materials form thioether bonds via step growth polymerization. This reaction is significantly faster and less toxic than chain-growth polymerization because it is not inhibited by oxygen and even consumes reactive oxygen species.^20,21^ Norbornene-modified biomaterials have been used successfully to encapsulate immortalized cell lines^21,36,37^ and human mesenchymal stem cells^20^ with biomimetic mechanical properties and may have potential for use with other primary cells as well. Furthermore, a unique requirement for on-chip culture of patterned features is their stability under continuous fluid flow so that patterned features are not dissolved or washed away. Without systematic testing of these factors, it remains difficult for laboratories to adopt on-chip photopatterning without extensive materials optimization.

In this work, we developed a straightforward protocol to pattern 3D cell cultures into customizable, free-standing structures under fluidic control, by using photolithography and photo-crosslinkable hydrogels within microfluidic devices. Using gelatin-based biomaterials with either methacryloyl or thiol-ene polymerization chemistries, we determined the effect of cell encapsulation on the storage moduli of each type of hydrogel and established conditions for biomimetic stiffness typical of soft tissues. We tested the effect of gelation chemistry and storage modulus on resolution and accuracy of the patterning, as well as the mechanical stability of free-standing micropatterns. Using this method, we demonstrated the capacity to create self-standing arrays and complex, non-linear hydrogel features using sequential photomasks. Finally, we demonstrated the use of in-situ photo-patterning for primary human cells, specifically T lymphocytes, and determined the cytocompatibility and spatial specificity of this micropatterning method.

## II. Results & Discussion

### A. Gelation chemistry and cell encapsulation

To develop a photopatterning method for 3D culture inside a microfluidic device, we started by identifying matrix components, gelation chemistry, and wavelength of light to support cell viability and function. For simplicity, we selected a gelatin backbone. As a naturally-derived material, gelatin provides cell adhesion (RGD) motifs and protease cleavage sites, eliminating the need to dope in other materials for this purpose.^38^ Inclusion of cells in the hydrogel precursor meant that the potential toxicity of both the cross-linking chemistry and the light exposure had to be minimized. Two gelatin-based biomaterials with different polymerization chemistries were tested: methacryloyl-modified gelatin (GelMA), which is widely used, under goes chain-growth polymerization and does not require a linker (**Fig. 1a**), and thiol-modified gelatin (GelSH) with a norbornene-linker, which undergoes biocompatible thiol-ene cross-linking chemistry (**Fig. 1b**). An 8-arm PEG-norbornene (PEG-NB) was selected as the linker; multi-arm linkers form stiffer hydrogel networks than dual-arm equivalents at a given dose of light exposure,^20^ which helped reduce exposure time and thus photo-toxicity. In the GelSH – PEG-NB system, both the protein and the PEG linkers were multifunctional, allowing cross-linking to occur during polymerization (**Fig. 1b**). GelSH is available commercially and is complimentary to norbornene-modified gelatin (GelNB) materials that have been described previously for 3D cell culture, but are not yet available with commercial quality control.^20,21,39^ For both materials, the photo-initiator lithium phenyl-2,4,6-trimethylbenzoylphosphinate (LAP) was chosen due to its ability to absorb violet-blue light (405 nm).^40^ This wavelength is less cytotoxic to cells than the UV light (365 – 385 nm) that is required by more common photoinitiators such as Irgacure-2959.

**Fig. 1.**
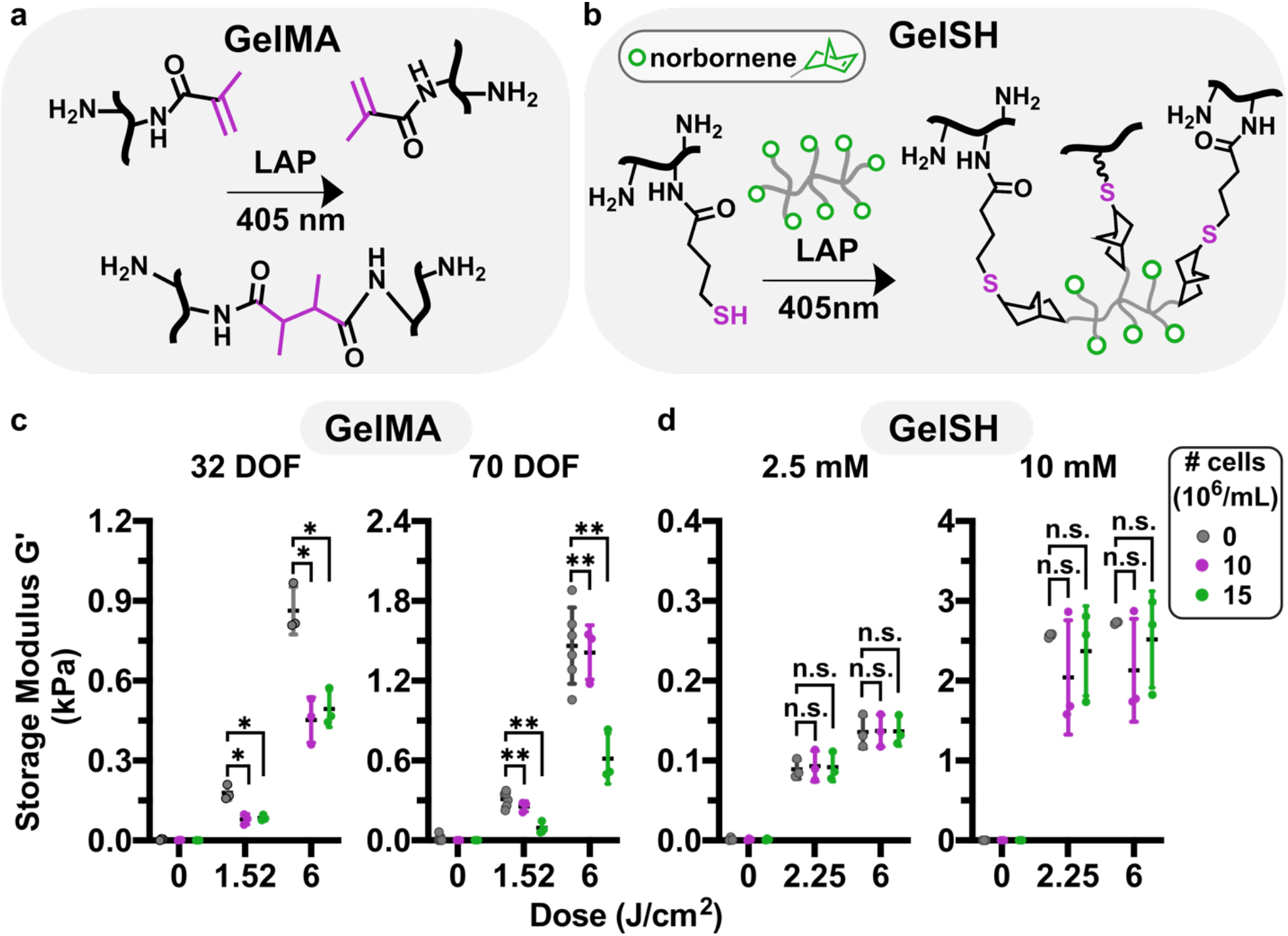
Impact of cell encapsulation on storage modulus of methacryloyl and thiol-ene hydrogels. (a) Reaction scheme of GelMA (black, reactive carbons in magenta) crosslinking in presence of LAP and 405 nm light. (b) Reaction scheme of GelSH (black, thiol in magenta) with an 8-arm PEG-norbornene linker (grey, NB in green), catalyzed by the photoinitiator LAP and 405 nm light. (c,d) Shear storage moduli of GelMA and GelSH hydrogels formed in the presence or absence of cells, at varying doses of light, for (c) GelMA (10% w/v) at 32 or 70 % DOF, and (d) 5% w/v GelSH cross-linked with 2.5 or 10 mM norbornene. Legend gives density of human CD4+ T cells, in 10^6^ cells/mL. n.s. p>0.05, * p≤0.05, ** p≤0.01 via Two-way ANOVA with Sidak’s multiple comparisons.

Since our approach to gelation required cells being present in the precursor as it polymerized, we investigated the extent of which the presence of cells at high densities influenced the mechanical properties of the various hydrogels. In initial experiments in the absence of cells, the concentrations of gelatin-based macromer and linkers were selected to provide a biomimetic range of shear storage moduli (G’), 120 – 3000 Pa. This range matches of that typical soft tissues such as brain^41^ and lymphoid tissue (**Fig. S1**).^42,43^ To access the lower and upper limits of the range, we used GelMA with varying degrees of functionalization (DOF) or varied the concentration of NB linker for GelSH, while the concentration of macromer was held constant. Inclusion of cells substantially altered the stiffness of GelMA hydrogels (**Fig. 1c**). At low DOF (32%), 10 to 15 × 10^6^ cells/mL decreased the shear storage modulus by approximately 2-fold. At high DOF (70%), 15 × 10^6^ cells/mL had a similar effect. These data suggested that high cell densities may hinder the methacryloyl groups from reacting with one another, particularly at lower DOF. In contrast, in the GelSH-NB system, inclusion of cells had no impact on the storage modulus at low (2.5 mM) or high (10 mM) concentrations of the NB linker (**Fig. 1d**). We speculate that the lack of effect of cells on storage moduli may be related to the network organization of the step-growth polymerized thiol-ene hydrogel. In the 10 mM NB hydrogels, G’ values varied by ~ 2-fold in the presence of cells, which we attribute to the challenge of adequately mixing the highly viscous precursor solution while maintaining cell integrity. While for some applications such variability may cause changes to biological activity, for our purposes the resulting G’ values were within the range of interest. In sum, these formulations of GelMA and GelSH were able to form soft (< 1 kPa) hydrogels in the presence of high densities of cells, but GelSH was less affected and retained the ability to form gels in the 1-3 kPa regime as well.

### B. Device Design and Micropatterning Process

Having established appropriate formulations for both types of photo-reactive hydrogels, we designed a simple and robust microfluidic housing for the patterned 3D cultures. The device was constructed from a thin layer of polydimethylsiloxane (PDMS) that was irreversibly bonded to a glass coverslip (**Fig. 2a**). PDMS is a well characterized, gas-permeable polymer that has been successfully used for many on-chip cell culture applications, is readily silanized to control surface chemistry, and is transparent for photo-crosslinking and optical imaging.^44^ For this work, the microchamber was designed with a 130-μm depth, sufficient to mimic a 3-dimensional tissue structure, and a 4.4-mm diameter at its widest point to provide sufficient surface area for complex patterns. These dimensions could be tailored readily in the future for specific applications. The chamber was tapered at each end to facilitate smooth filling.^45^ In order to covalently anchor the patterned, norbornene- or methacryloyl-bearing hydrogel to the surface of the chip, the interior of the device was oxidized in a plasma cleaner and functionalized with either a thiol-or methacrylate-terminated silane, respectively (**Fig. 2b**). It was critical not to over-treat the PDMS surface with oxygen plasma, which caused cracks in the PDMS that templated defects in the hydrogel (**Fig. S2**).^46^

**Fig. 2.**
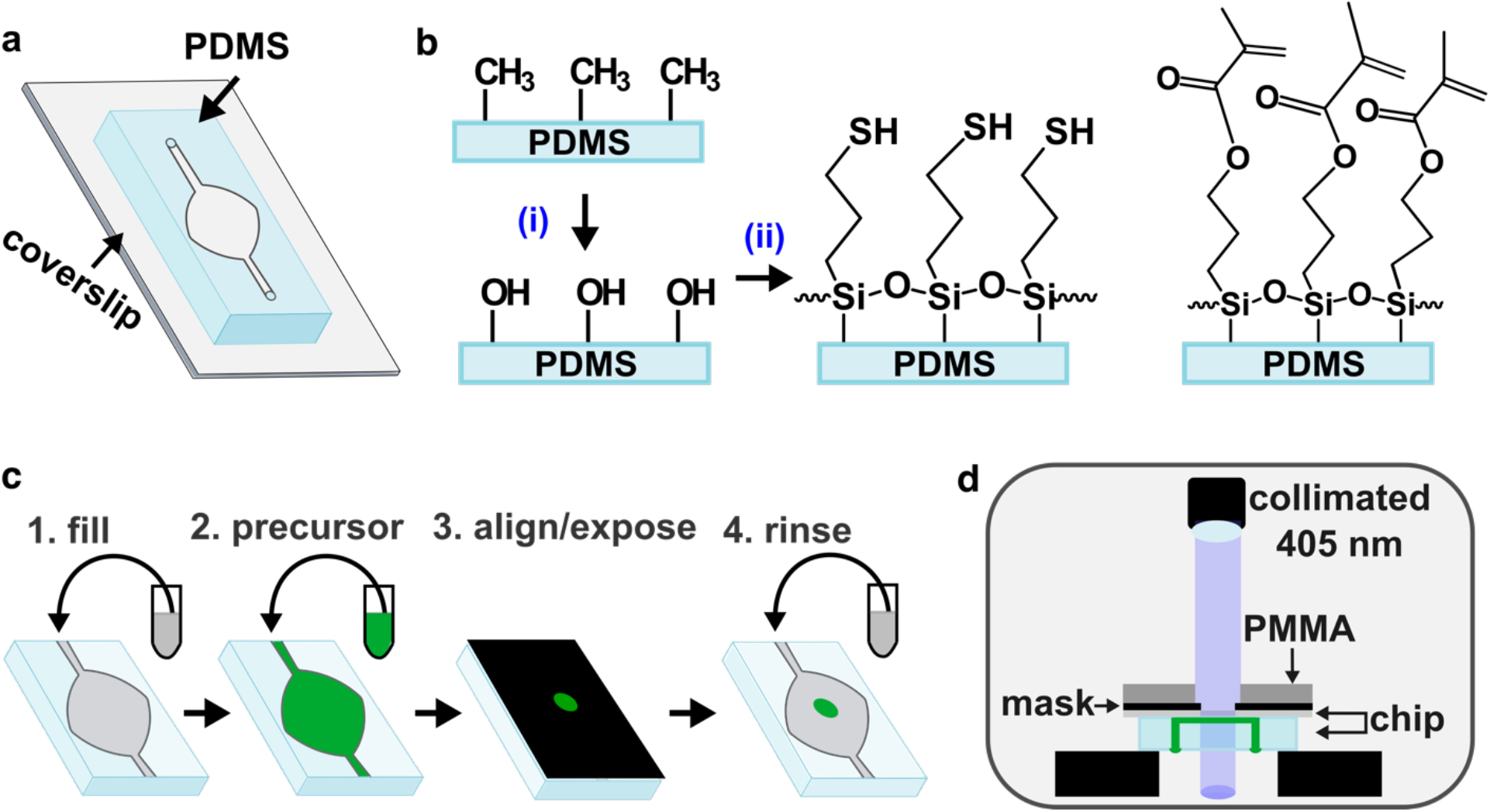
Photo-patterning set-up and process. (a) Schematic of chip: thin layer of PDMS bonded to a glass coverslip. (b) Surface functionalization of PDMS. The methyl surface was (i) activated via oxidation with air plasma, followed by (ii) silanization using either thiol-terminated (left) or methacrylate-terminated (right) silane, to match the intended hydrogel. (c) Stepwise schematic of patterning process: 1) The chip was filled with buffer (grey), and 2) the buffer was displaced by precursor (green). 3) A photomask with desired design was aligned against the coverslip, supported with a rigid polymer backing (PMMA), clamped (not shown for clarity), and exposed. 4) Unreacted material was removed with a buffer rinse. If needed, the process was repeated with a different precursor to add additional structures. (d) Schematic of photo-patterning set up. The chip was placed upside down on top of two support layers (black) to suspend it below the collimated light source. The channels and chamber of the chip are shown filled with precursor (green).

The patterning process consisted of four steps (**Fig. 2c**): After sterilization, the device was pre-wet with phosphate-buffered saline (PBS), filled with a precursor solution, and sealed off to prevent air entry. Precursor solutions consisted of gelatin monomer and any linkers, LAP, and the desired population of cells in suspension. The chip was placed briefly on a cooling stage to lower the temperature and ensure consistent gelation. Next, the photo-mask was aligned against the coverslip of the chip (**Fig. S3**), and the chip was placed upside down below a collimated 405 nm LED light source and exposed (**Fig. 2d**). Finally, the inlet was unplugged, and unreacted precursor was removed by flowing PBS, leaving only the patterned culture. The process was sequentially repeated with additional precursor solutions, e.g., containing different material compositions or different populations of cells. Once all patterning steps were completed, culture media was flowed into the chip and transferred to a cell culture incubator for continued culture.

### C. Resolution, robustness and stability of on-chip photolithography

The resolution and fidelity of patterning thick hydrogels by on-chip photolithography is expected to be limited by scattering of the incoming light and by diffusion of reactive species, as well as the mechanical properties of the hydrogel itself.^34^ Therefore, we tested the resolution and ability to pattern GelMA and GelSH hydrogels, whose different mechanisms of polymerization result in different organization of crosslinked networks.^19^ We initially hypothesized that hydrogels with a lower storage modulus, due to their lower cross-linking density, would have higher swelling ratios,^47^ which would result in dimensions larger than intended. Resolution and accuracy of on-chip patterning were tested by using photomasks with circular features ranging from 100 to 900 μm in diameter (**Fig. 3a**). Chips were filled with precursor, exposed, rinsed with PBS to remove un-crosslinked material, and imaged immediately to determine the dimensions of the freshly patterned structures (**Fig. S4**). To remove any poorly crosslinked regions, chips were imaged again to determine the dimensions of the features after a 30-min incubation and rinse (“incubated”; **Fig. 3b**). While circular free-standing features are very difficult to achieve on microfluidic chips with standard methods, they were straightforward to produce by on-chip photolithography.

**Fig. 3.**
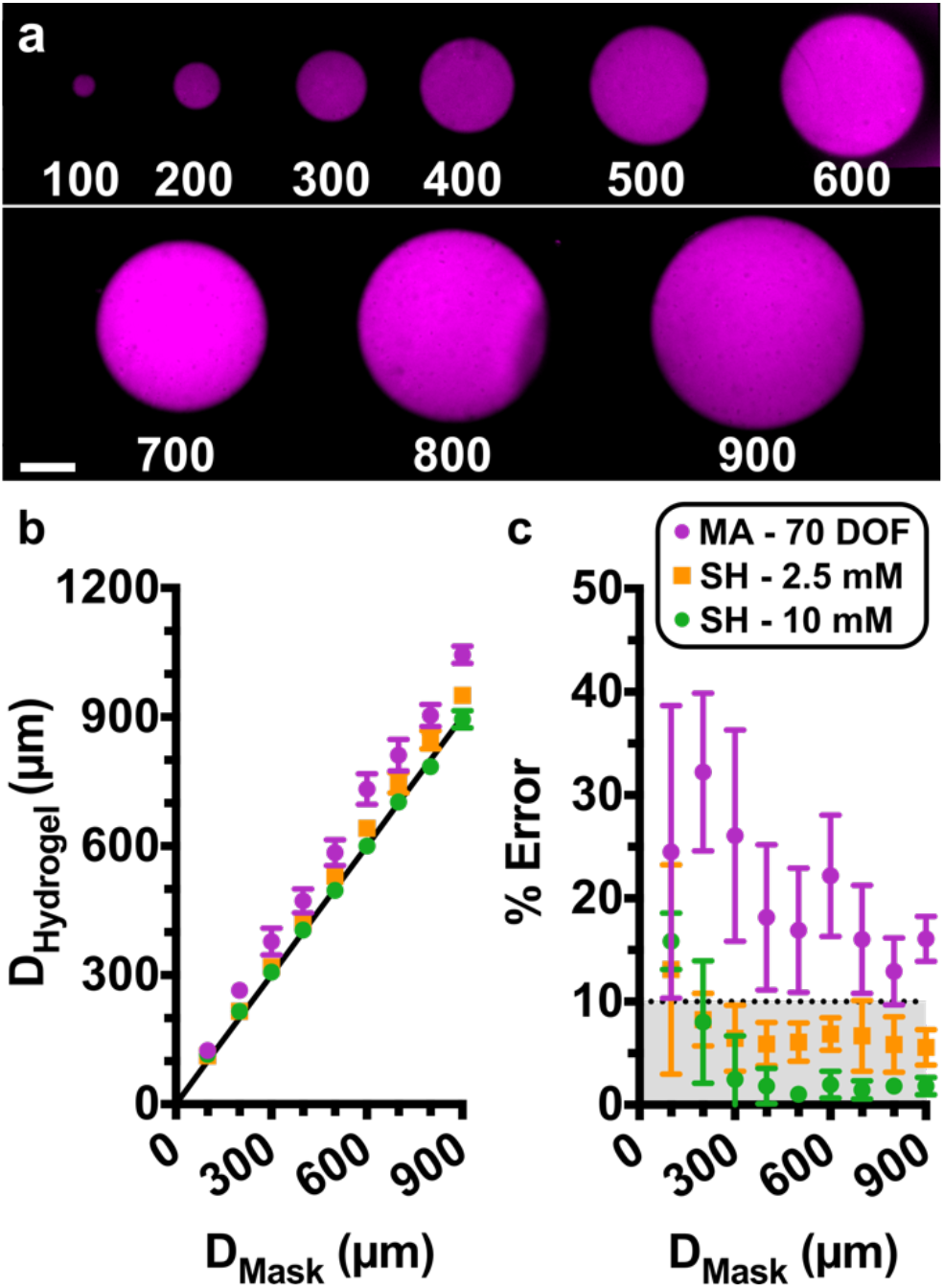
Assessing Pattern Resolution. (a) Fluorescent images of circular hydrogel features patterned on a microfluidic chip. Features ranged from 100-900 μm in diameter. Shown are features in GelSH with 2.5 mM NB, labelled with NHS-rhodamine, on a chip with a 0.15-mm coverslip. Scalebar 250 μm. (b-c) Quantification of accuracy. (b) Plot of measured diameter of the hydrogel region versus the diameter of the design on the photomask. Black line represents y = x, shown for reference. Measurements were taken after a 30-min incubation and rinse. (c) Calculated percent error of each feature versus the target diameter from the photomask design. The dotted line was drawn arbitrarily at 10% error, and grey area shows values that fall in that region. The shared legend shows 10% GelMA with 70% DOF (n=3) and 5% GelSH with norbornene concentrations of 2.5 mM (n=4) or 10 mM (n=4). Symbols and error bars represent mean and standard deviation; some error bars too small to see.

As expected, all hydrogel features were linearly dependent on the dimensions defined by the photomasks (**Fig. 3b**). Furthermore, features were obtained reproducibly down to 100 μm, the smallest size tested, for GelSH and for 70% DOF GelMA. In stiffer GelSH hydrogels (G’ > 1 kPa, 10 mM NB), the feature dimensions obtained were highly accurate, matching those of the photomask with < 10% error for all feature sizes, except for 100 μm which had 16% error (**Fig. 3c**). Features patterned with lower storage moduli (G’ < 0.5 kPa, 2.5 mM NB) were slightly larger than intended, but nevertheless also had < 10% error when larger than 100 μm (**Fig. 3c**). For both GelSH materials, there was no significant change in dimensions between freshly patterned and incubated features, suggesting all weakly cross-linked or un-crosslinked material was fully removed during the first rinsing step, and that no significant swelling took place during incubation (**Fig. S4).** On the other hand, in stiffer GelMA hydrogels (G’ > 1 kPa, 70% DOF), the hydrogel features were larger than the dimensions of the photomask by 13 – 32%. The larger features may be a result of the relatively long exposure time needed to generate the desired storage modulus, which may allow for diffusion of reaction species beyond the area illuminated by the photomask. Furthermore, the GelMA features grew significantly after incubation (**Fig. S4**), suggesting significant levels of swelling. Features formed using soft GelMA (G < 0.5 kPa, 32% DOF) were observable after initial exposure but dissolved completely after a 30 min incubation period (not shown), making this formulation unsuitable for patterning. Therefore, the accuracy of patterning and initial feature stability was dependent not just on storage modulus, but also on the chemistry of gelation.

While the data above were collected using a glass coverslip (0.15 mm) for the bottom of the device, we also tested the extent to which resolution of features in GelSH was affected by the use of a thicker glass layer (1 mm), which is often preferred over coverslips to make more robust chips. Under this condition, while the diameter of the patterned features remained linearly dependent on the dimensions of the photomask, a small corona was formed around the patterns, resulting in larger dimensions (**Fig. S4**). The corona largely disappeared following incubation, indicating the formation of loosely cross-linked materials that later dissolved or were rinsed out. With the 1 mm glass, features were obtained reproducibly down to 200 μm in all tests; 100 μm features did not gel consistently. Therefore, while better resolution and pattern fidelity were obtained with the 0.15 mm coverslip, the more robust chip may be an acceptable tradeoff for applications where larger features are sufficient.

Finally, we tested the robustness of the patterning method to a change in light source. As expected, reproducible gelation absolutely required the use of the collimated light source. Patterning with uncollimated light under the same conditions resulted in inconsistent gelation across chips, in some cases with regions left un-crosslinked, or with features that were weakly cross-linked but washed out after the 30-min incubation period (data not shown). This phenomenon was consistent with the need for collimated light to provide uniform light intensity during photolithography in other settings, i.e., to mitigate light scattering, interference, and heterogenous dose across the exposed area. ^48,49^

### D. Micropatterned feature stability under overnight perfusion

Having established the resolution and stability of patterned features under static conditions, we next assessed the mechanical stability of the patterned hydrogels under conditions mimicking those required for cell culture, i.e., under continuous perfusion overnight inside a humidified cell culture incubator (**Table 1**). GelSH hydrogels of higher storage modulus (10 mM NB) maintained 100% of the features that were 300 μm and larger, while smaller features, 200 and 100 μm, were stable in 75% and 50% of chips, respectively. For GelSH hydrogels of lower storage modulus (2.5 mM NB), we observed that features larger than 300 μm remained stable in majority of the chips patterned (75%), while features smaller than 300 μm were unstable in all chips. We speculate that the greater stability of larger GelSH features may be due to their higher contact area and thus more crosslinks to the PDMS and glass surfaces of the chip, or to their smaller surface/volume ratio that reduces exposure to shear flow in the chamber. In contrast to the relative stability of the GelSH features, even the stiffer GelMA hydrogels (70% DOF) proved unstable after overnight perfusion in features of all sizes. 32% DOF GelMA hydrogels were not tested under these conditions, because features were not stable after a 30 min incubation period.

**Table 1.** Stability of patterned hydrogel after overnight perfusion. Reported values are % of successful attempts of features that remained anchored on the chip, as opposed to dissolved or rinsed away, after overnight perfusion. N=4 chips for all hydrogel formulations. (*) indicates not tested due to feature instability at short times.

| Macromer | Formulation | Expected G' (kPa) | Feature Size (μm) |  |  |  |  |  |
| --- | --- | --- | --- | --- | --- | --- | --- | --- |
|  |  |  | 600 | 500 | 400 | 300 | 200 | 100 |
| 5% GelSH | [NB] |  |  |  |  |  |  |  |
|  | 10 mM | 1 | 100% | 100% | 100% | 100% | 75% | 50% |
|  | 2.5 mM | 0.1 | 75% | 75% | 75% | 75% | 0% | 0% |
|  | % DOF |  |  |  |  |  |  |  |
| 10% GelMA | 70 | 1 | 0% | 0% | 0% | 0% | 0% | 0% |
|  | 32 | 0.1 | * | * | * | * | * | * |

These data again revealed differences in feature stability as a function of both the storage modulus and the gelation chemistry. This may ultimately be a consequence of the cross-linking density of the network, with less dense networks being less mechanically stable. Indeed, for both chemistries, the micropatterned features were less stable in softer gels, which have fewer crosslinks than stiffer gels. Furthermore, under these conditions, micropatterned GelSH hydrogels proved to be more physically stable than micropatterned GelMA hydrogels, regardless of storage modulus. We speculate that the instability of GelMA features may be related to the short exposure times that were required to minimize cytotoxicity, which constrained the material to the linear stage the polymerization reaction, where cross-linking is incomplete (**Fig. S1**). On the other hand, the faster thiol-ene polymerization reached a saturated cross-linking density in this time period. We note that physical stability of patterned features also likely depends in part on the quality of the surface functionalization with reactive silane; indeed, we observed changes in overnight stability when a different plasma cleaner was used. In summary, the stability of micropatterned features under flow was a function of hydrogel formulation, storage modulus (likely as a proxy for degree of crosslinking), and the surface functionalization of the chip. Based on the high accuracy and stability observed in GelSH hydrogels, as well as published reports of biocompatibility,^20,21^ we utilized GelSH for all subsequent experiments.

### E. Complex geometries via multi-step photopatterning of GelSH hydrogels

Working with the thiol-norbornene hydrogel system, we tested the extent to which on-chip photolithography provided access to micropatterned hydrogel geometries at increasing levels of complexity. These experiments were intended to test the patterning system’s ability to generate various types of features of interest in future experiments, rather than to test particular biological functions. First, we tested the ability to pattern open channels and a curved fluidic path (**Fig. 4a**), which will be critical for future use in patterning vascularized systems or multiple stand-alone culture regions. By preventing exposure of the precursor solution in the center and near the walls of the chamber, we were able to pattern two self-standing lobes divided by an open, central channel through which fluid could flow. Next, we tested geometries that are challenging to achieve by standard microfluidic patterning methods. It was straightforward to pattern regions with shared, non-linear boundaries, e.g., by creating a self-standing island followed by a second surrounding hydrogel (**Fig. 4b**). The two regions were visually in contact under microscopic imaging, without a gap. We extended this system to pattern three sequential regions in concentric circles (**Fig. 4c**). Nonlinear adjacent regions will be useful in the future for cellular invasion assays, angiogenesis assays, and patterning of biomimetic tissue structures in the future. Finally, to test the versatility with more intricate geometries and alignment capabilities, we recreated the University of Virginia (UVA) historic Rotunda by patterning hydrogels in three sequential steps: 1) the columns, 2) the negative space surrounding the columns, and 3) the dome and foundation (**Fig. 4d**). In all cases, the second and subsequent patterns were achieved through the use of photomasks that covered the previously patterned constructs, ensuring that each region only received one dose of light. We note that these experiments used the same parent microfluidic chamber for all designs; only the photomasks were changed. Thus, the spatial organization of the patterned gel was altered rapidly between subsequent devices, without time-consuming master fabrication.

**Fig. 4.**
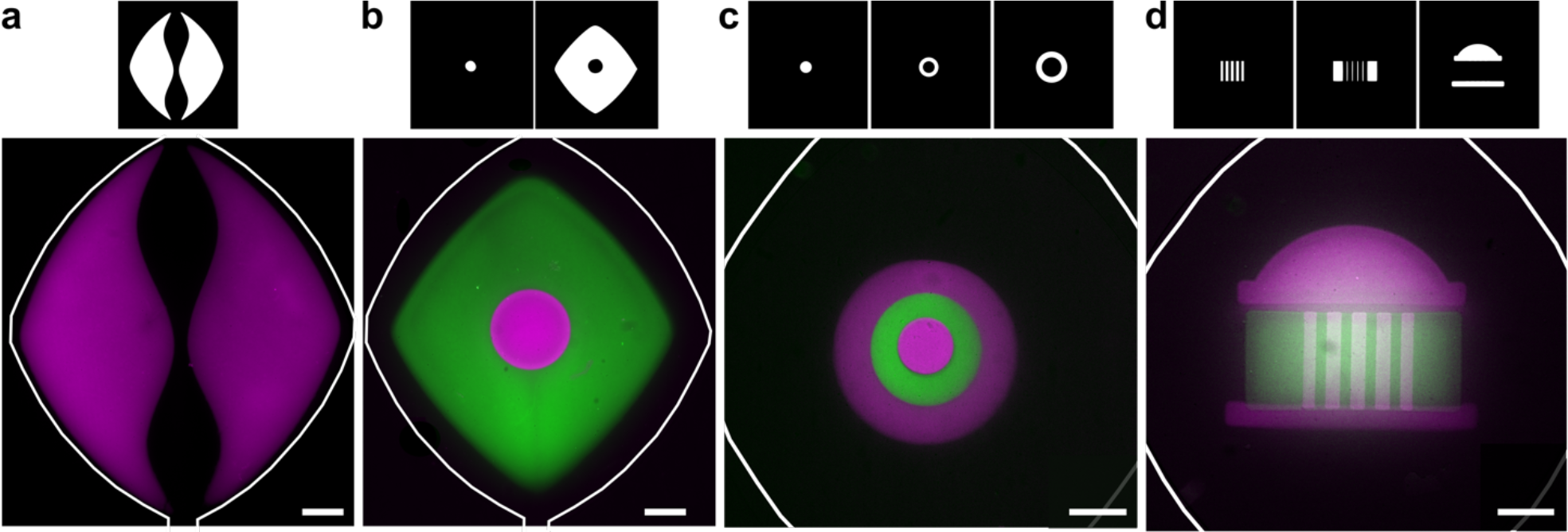
Geometric versatility achieved by on-chip photo-patterning of GelSH hydrogels. (a) NHS-rhodamine-labelled hydrogel (magenta) used to pattern a curved fluidic path in culture chamber. (b) A central circular island (magenta) surrounded by NHS-fluorescein-labelled GelSH (green). (c) Concentric circles patterned with hydrogel labelled with NHS-rhodamine and NHS-fluorescein in three sequential steps. (d) A patterned UVA Rotunda in three sequential steps. The corresponding photomasks used to achieve patterns are shown above each panel. All scalebars are 500 μm.

### F. Micropatterning of cell-laden hydrogel features on chip

Next, we tested the ability to pattern cell-laden features in targeted locations on chip. Primary naïve human T cells (CD4+) were used as a rigorous case study; these non-proliferative cells are of interest for on-chip testing of immunotherapies. The T cells were suspended in the precursor solution immediately before loading it onto the chip for patterning. As with the cell-free patterns, the cell-laden un-crosslinked hydrogel-precursor was readily washed out from designated regions in the center and edges of the chamber, to generate open channels (**Fig. 5a,b**). Next, we tested the ability to pattern complex, cell-laden geometries with a stylized alien facial pattern. The central eyes and mouth were patterned first, followed by the surrounding head shape (**Fig. 5c,d**). The resulting features were composed of two separate cell populations that shared non-linear boundaries with one another. These geometries would be challenging to obtain on-chip by laminar flow or by surface-tension, even with the inclusion of micropillars.

**Fig. 5.**
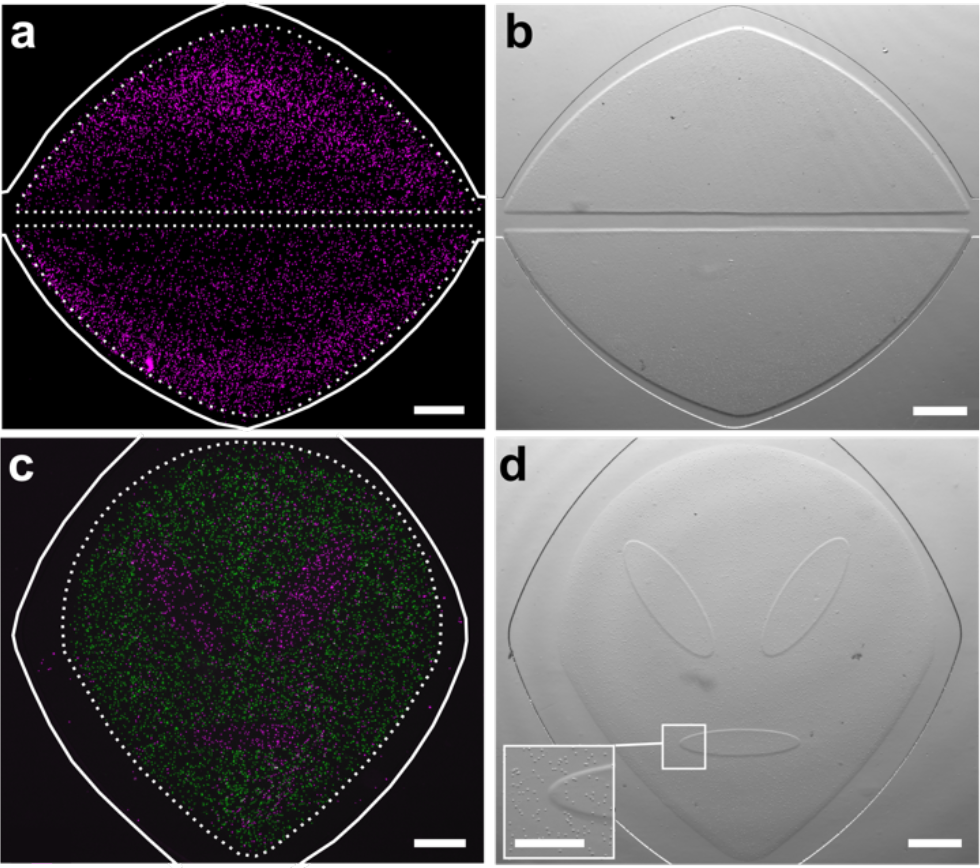
*In situ* photo-patterned cell-laden hydrogel constructs. (a) Fluorescence and (b) brightfield images of a patterned 3D cell culture (cells labelled magenta), patterned into two self-standing lobes. A linear fluidic path was patterned between them, and a second, curved fluidic path surrounded them for better distribution of media. (c) Fluorescence and (d) brightfield images of two distinct cell populations patterned into an alien face geometry. First cell population labelled with NHS-rhodamine (magenta); second population labelled with CFSE (green). Inset shows magnified boundary between two patterned regions. Scale bar is 500 μm in (a-d), 250 μm in inset. Dashed lines denote the boundary of the hydrogel regions; solid white lines indicate the edges of the microfluidic culture chamber.

We note that while there is great flexibility to the types of geometries that can be achieved with this method, one is limited by the requirement to rinse out un-crosslinked materials. In particular, concave structures and shapes with voids, such as the letters A and O, are not directly accessible, but multi-step patterning offers a potential solution to this issue. For example, to pattern a cell-laden ring around a cell-free center, the inner region would be patterned first using gel without cells, followed by the surrounding ring.

### G. Precise patterning and viability of primary CD4 T cells in GelSH microarray on chip

Next, we investigated the spatial precision of cellular patterning and whether it was dependent on hydrogel formulation. During the loading of the chip, the cell-laden hydrogel-precursor fills the entire culture chamber, giving cells an opportunity to non-specifically adhere to the surfaces of the chip outside the intended patterned regions. To rigorously quantify the specificity of cell location in the patterns, we created an array of 9 circular features per chip in diameters of 200, 400, and 600 μm. These dimensions are representative of the length scale of tissue substructures in complex organs like brain, lymph nodes, and solid tumors.^50,51^ Cells were patterned at high density (> 10^7^ cells/mL) in GelSH hydrogels with 2.5- or 10-mM NB linker (**Fig. 6a,b**), and incubated under continuous fluid flow overnight. As expected, the mean density per unit area in the patterned regions was high (**Fig. 6c**), and feature size had no effect on cell density (data not shown). Non-specific adhesion was minimal outside of patterned hydrogels, as the cell density in the non-exposed regions was less than 4.5% of that in the patterned areas (**Fig. 6c**). The high efficiency of targeted patterning may be the result of the rapid precursor loading and short exposure times, which allowed for the rinsing step to start less than one minute after cells enter the chamber. Thus, cells were patterned precisely in the intended regions, with minimal adhesion elsewhere.

**Fig. 6.**
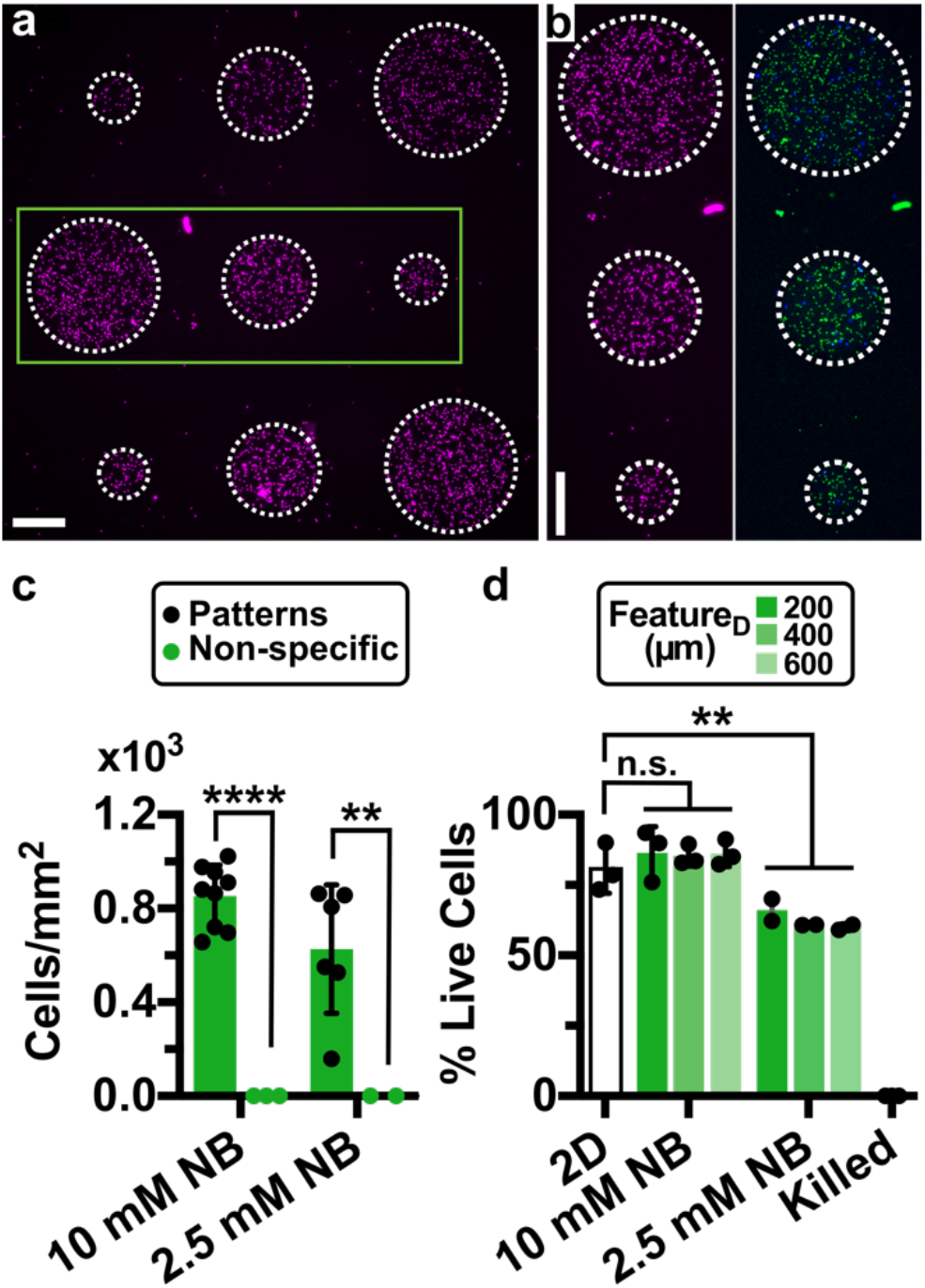
Precision and viability of photo-patterned microarray of human CD4 T cells on chip. (a) Nine-circle culture array patterned on-chip with cells pre-labelled with NHS-rhodamine. Scalebar 250 μm. (b) Zoomed-in view of area outlined in green in panel 6a. (*Left)* Image of NHS-rhodamine labelled cells; (*Right*) image after viability staining with Calcein-AM (green) and DAPI (blue). Scalebar 250 μm. (c) Quantification of cell density inside and outside of the patterned regions in GelSH hydrogels (n=2 and n=3 chips respectively). Two-way ANOVA with Sidak’s multiple comparisons; **** p≤0.0001, ** p≤0.01. (d) Quantification of the viability of patterned CD4+ T cells in 5% GelSH with 2.5 and 10 mM NB hydrogels as a function of feature dimensions after overnight culture under continuous fluid flow, versus off-chip (2D) controls (n.s. p > 0.05, ** p≤0.01 One-way ANOVA with Tukey’s multiple comparisons, n=3 and n=2 for 10 mM and 2.5 mM NB chips, respectively).

Finally, the micropatterned GelSH-based culture arrays were cultured overnight to test the initial effects of patterning and pattern geometry on overnight survival of these fragile primary cells. Cultures were held under continuous flow of media to ensure replenishment of nutrients and oxygen in the microfluidic chip. Using the microfluidic culture system, it was straightforward to deliver staining reagents at the end of the experiment to measure viability in situ by flowing in a Calcein-AM/DAPI (live/dead) solution, incubating, rinsing, and imaging (**Fig. 6b**). In 10 mM NB hydrogels, there was no significant difference in the percentage of live cells between on-chip cultures and off-chip unpatterned controls (**Fig. 6d**) nor between the feature dimensions. Interestingly, viability was slightly reduced in the 2.5 mM NB hydrogels compared to off chip controls (20% decrease, p=0.0089, One-way ANOVA with Tukey’s multiple comparisons), though still in an acceptable range. As the focus of this work was on the development of the micropatterning method, we did not further explore the impact of gel chemistry and the internal structure of the hydrogel on longer term cell viability and behavior; these will be exciting areas for future investigation.^52–55^ These experiments confirm that the on-chip photopatterning method was cytocompatible and ready for future implementation to study the impact of spatial organization on cell function, including with primary cells.

## III. Conclusion

In summary, we have described a protocol for in situ micropatterning of spatially organized biomaterials and 3D cell cultures on a microfluidic chip and established the impact of crosslinking chemistry on the storage modulus, stability, and spatial resolution of the patterns. By simply aligning a photomask prior to light exposure, the user may pattern a wide variety of design configurations in the xy-plane without altering the microfluidic housing. The resulting patterned cultures were modular and free-standing, without the need for physical supports such as micropillars to guide the hydrogel in place. Gelation chemistry had a significant impact on the accuracy and mechanical stability of patterned microfeatures. While features were patterned down to 100 μm in both GelSH and GelMA hydrogels, the GelSH hydrogels in stiffer formulations provided the highest accuracy and greater stability under fluid flow. Complex geometries such as concentric circles, architectural designs, and microarrays were all accessible, as were open flow paths to distribute media to the patterned 3D cultures. When used with thiol-ene polymerization chemistry, the micropatterning method had high specificity and low cytotoxicity with primary human cells. We envision that this micropatterning strategy will enable researchers to organize 3D cultures directly onto microfluidic chips in arrangements that capture the complexity of tissue organization, thus granting access to mechanistic experiments while maintaining control over cellular and fluidic components.

## IV. MATERIALS & METHODS

### A. Hydrogel materials and sourcing

Thiol-modified gelatin (GelSH; Lot: MKCJ5413) was obtained from Sigma Aldrich and used as provided. The vendor-reported absolute degree of functionalization for this material, determined by free thiol assay, was 0.223 mmol -SH / g gelatin. 8-arm PEG-NB 20 kDa (Jenkem Technologies), lithium phenyl-2,4,6-trimethylbenzoylphosphinate (LAP; Sigma Aldrich), and 1x phosphate buffered saline without calcium or magnesium (1x PBS; Lonza) were also used as provided. Gelatin methacryloyl (GelMA; Lots: MKCK4076 and MKCK5644) with vendor-reported fractional degrees of functionalization of 70% and 32%, respectively, were obtained from Sigma Aldrich. The absolute degrees of functionalization were also measured in-house by H-NMR, as described by Zatorski et. al., (2020),^56^ and found to be 0.232 and 0.088 mmol -MA/g GelMA, respectively.

### B. Sourcing of human CD4+ T cells

Human naïve CD4+ T cells were purified from TRIMA collars, a byproduct of platelet apheresis, obtained from healthy donors (Crimson Core, Brigham and Women’s Hospital, Boston, MA and INOVA Laboratories, Sterling, VA). Initially, total CD4+ T cells were isolated using a combination of the human CD4+ T cell RosetteSep™ kit (STEMCELL Technologies) and Ficoll-Paque (Cytiva Inc.) density centrifugation. Naïve CD4+ T cells were then enriched from total CD4+ T cells through immuno-magnetic negative selection with the EasySep™ Naïve CD4+ T cell isolation kit (STEMCELL Technologies). Naïve CD4+ T cell post-isolation purity (CD4+CD45RA+CD45RO-) was determined through flow cytometry (**Fig. S5**).

### C. GelSH and GelMA Matrix Characterization and Cell Encapsulation

For rheological measurements of GelSH hydrogels, the precursor solution was prepared by combining reagents to a final concentration of 5% w/v GelSH, 2.5 or 10 mM norbornene (0.313- or 1.25-mM PEG-NB linker), and 3.4 mM LAP in 1x PBS. The PEG-NB linker was added right before the sample was to be pipetted onto the stage of the rheometer. For GelMA, reagents were combined to a final concentration of 5 or 10% w/v GelMA and 3.4 mM LAP in 1x PBS. When cells were included during photopolymerization, cells were spun down and resuspended at 10 or 15 × 10^6^ cells/mL in precursor solution.

Rheological characterization was performed using a MCT302 Anton Parr Rheometer, operated in oscillatory time sweep mode with 5% strain, 1 Hz frequency, and 0.1 mm gap to assess gel polymerization rate and storage modulus. A UV-curing stage was fitted with a 20-mm parallel plate, the light source was filtered through a 400-500 nm filter, and the stage temperature was maintained at 25 °C. 30 μL of precursor solution was pipetted onto the stage. After measuring baseline shear storage modulus for 30 seconds, light exposure was initiated with constant intensity of 50 mW/cm^2^ and continued for 2 min.

### D. Microfluidic Device Fabrication and Surface Modification

A one-layer microfluidic device was fabricated using standard soft lithography methods. Transparency masks were drawn in AutoCAD LT 2019 and printed at 20,000 DPI by CAD/Art Services, Inc. (Brandon, OR). The master molds were fabricated using SU-8 3050 photoresist spun to 124 – 136 μm thickness (Microchem, Westborough MA, USA) on 3” silicon wafers (University Wafer, South Boston MA, USA) and vapor silanized with Trichloro(1*H*,1*H*,2*H*,2*H*-perfluorooctyl) silane (Sigma Aldrich) for 2 hours. Degassed polydimethylsiloxane (PDMS) was prepared at a 10:1 ratio of elastomer base to curing agent (Slygard 184 Silicone Elastomer, Ellsworth Adhesives, Germantown WI, USA), poured over the silicon SU-8 master, and cured in a 70 °C oven for at least 2 hours.

Once cured, the PDMS was removed from the master and punched at the channel ends using a 0.75-mm I.D. tissue punch (World Precision Instruments, Sarasota FL, USA) to create inlets for PFTE TT-30 tubing (Weico Wire Inc.). The PDMS layer and a Goldseal cover glass (35 × 50 mm × 0.15 mm, actual thickness 0.13-0.16 mm, Ted Pella, Inc.) were oxidized in a plasma cleaner for 20 seconds (air plasma; Tegal Plasmod) or for 10 seconds with BD-20AC laboratory corona treater^57^ (Electro-Technic Products, Chicago IL, USA), manually assembled, and incubated in a 120 °C oven for 10 minutes to complete the bonding process. Where noted, ~ 1 mm thick Corning^®^ Glass Slides, 75 × 50 mm (Ted Pella, Inc.) was used instead of the glass coverslip. The device was then purged with nitrogen for 10 min, followed by 90 min of vapor silanization using (3-Mercaptopropyl) trimethoxy silane (Sigma Aldrich) in a nitrogen-filled environment. For devices to be used with GelMA, 3-(Trimethoxysilyl)propyl methacrylate (Sigma Aldrich) was used instead. After silanization, the device was rinsed with 70% ethanol and distilled water, purged with a nitrogen gun to remove excess moisture, and placed in a 120 °C oven to dry completely for at least 10 min. Once dried, the device was covered with tape to prevent dust accumulation and stored in a desiccator containing Dri-rite. SH-functionalized devices were used within a week of and MA-functionalized devices were used within 2 days of being silanized.

A rigid poly (methyl methacrylate) (PMMA) cover was fabricated to serve as a backing for the photomask. To avoid light scattering or absorption during photo-crosslinking, a hole was cut into the PMMA over the culture chamber. Specifically, a 50 × 45 × 1.5 mm acrylic sheet (McMaster-Carr, Princeton, NJ USA) was etched with a 10-mm central hole by using a CO_2_ laser (Versa Laser 3.5, Universal Laser System, Scottsdale, AZ) set to 20% power and 1% speed.

### E. Photo-patterning of spatially organized hydrogels and 3D cultures on chip

Prior to photopatterning, a precursor solution was prepared by combining reagents to a final concentration of 5% GelSH and 3.4 mM LAP in 1x PBS. This solution was incubated at 4 °C overnight, then warmed and kept at 37 °C during use. In some cases, the GelSH was fluorescently labelled prior to gelation by pipetting a reactive dye into the precursor solution and vortexing for 30 seconds at room temperature. Final concentrations of reactive dye were 5 μM NHS-rhodamine, 53 μM NHS-fluorescein, or 31 μM Alexa Fluor 594-succinimidyl ester (ThermoFisher). For experiments with cells, the cells were centrifuged at 400 × g for 5 min to remove culture media, resuspended in 1x PBS, and centrifuged again to remove 1x PBS immediately before resuspending in precursor solution. PEG-NB linker was added and carefully mixed immediately before loading the precursor solution into syringes.

The microfluidic chip was sterilized with 70% ethanol for 2.5 min, followed by a 5 min rinse with DI water and a subsequent 5 min rinse with 1x PBS, all at 5 μL/min. All rinse steps were performed by using a Chemyx syringe pump (Fusion 200, Houston TX, USA) and a 1-mL Hamilton™ Gastight Microliter™ syringe. Precursor solution was loaded into a 25-μL Hamilton™ 700/1700 Series Gastight™ syringe connected to TT-30 tubing (Weico Wire) and was flowed into the chip for 2 min at 5 μL/min. The inlets and outlets were blocked off with PDMS-filled TT-30 tubing to prevent air entry. Next, the chip was placed on an aluminum cooling plate with a reusable ice pack (Dulytek, Amazon) at 14 – 18 °C for 45 seconds to cool the device; this step ensured a consistent temperature across chips during gelation. The photo-mask was taped to the PMMA cover, aligned to the coverslip of the chip, and clamped using binder clips. The chip was then overturned and exposed to light using a Fiber Coupled Violet LED light source at 405 nm (±5 nm) attached to a Ø0.5-inch fiber collimator (Prizmatix, Inc., Israel). The light intensity was kept constant at 50 mW/cm^2^. The chip exposed for 45 sec to achieve the optimized dose of 2.25 J/cm^2^, unless otherwise noted. Once patterned, the device was returned to its upright position and rinsed with room temp 1x PBS for 5 min at 5 μL/min to remove uncrosslinked material. If additional patterning steps were necessary, device was re-loaded with a new precursor solution, cooled, aligned, exposed, and rinsed. Finally, for experiments in which cells were present, appropriate culture media was flowed in for 2 min at 5 μL/min before the chip was connected to perfusion as described below.

### F. Widefield Imaging

Except where noted below, imaging was performed on an upright Zeiss AxioZoom macroscope equipped with a HXP 200C metal halide lamp, PlanNeoFluor Z 1x objective (0.25 NA, FWD 56 mm), and Axiocam 506 mono camera. For fluorescence imaging, filters used were Zeiss Filter Set 38 HE (Ex: 470/40, Em: 525/50), 43 HE (Ex: 550/25, Em: 605/70); 64 HE (Ex: 587/25, Em: 647/70); and 49 HE (Ex: 365, Em: 445/50). Brightfield images were collected using transmitted light. Zen 2/3 Blue software was used for image collection, and images were analyzed in ImageJ v1.52k.

For imaging cells after overnight culture, we used a Zeiss AxioObserver 7 inverted microscope equipped with a Colibri.7 LED light source, EC Plan-Neofluar 5x objective (N.A.=0.16, WD=18.5 mm), and ORCA-Flash4.0 LT+ sCMOS camera (Hamamatsu). For fluorescence imaging, the filter used was a Zeiss 112 HE LED penta-band. Zen 3 Blue software was used for image collection.

### G. Assessing Patterning Resolution

Chips were assembled as described above, with the bottom layer comprised of either a coverslip (0.13 – 0.16 mm thickness) or a 1-mm thick Corning^®^ Glass Slides, 75 × 50 mm (Ted Pella, Inc.). In the case of GelSH, a precursor solution composed of 5% GelSH, 0.313- or 1.25-mM PEG-NB linker, 3.4 mM LAP, and 5 μM NHS-Rhodamine was patterned on-chip as described above, using a 45 second exposure at 50 mW/cm^2^. In the case of GelMA, a precursor solution composed of 10% GelMA, 3.4 mM LAP, and 5 μM NHS-Rhodamine was patterned on-chip as described above, using exposure at 50 mW/cm^2^. The exposure times were 119 and 30 seconds for 70% and 32% DOF GelMA, respectively. For GelMA samples, the chips were placed in the incubator immediately for 1 min after light exposure to reduce the viscosity of the material for rinsing. Un-crosslinked material was rinsed out with 1x PBS for 5 min at 5 μL/min, after which the inlet and outlet were closed using TT-30 (Weico Wire) tubing filled with PDMS. The chips were placed in an incubator (37 °C, 5% CO_2_) for 30 minutes in the absence of flow, then chips rinsed once more with 1x PBS for 5 minutes at 5 μL/min. Features were imaged by brightfield and fluorescence microscopy on a Zeiss AxioZoom microscope (HE 43 filter set). The diameter of each feature was quantified from the fluorescence images by using line tools in ImageJ v1.52k.

### H. Overnight perfusion of micropatterned features

To minimize accumulation of bubbles in the microdevice, home-made PDMS bubble traps were used based on the design described by Jiang et al.^58^ In short, a thick (3.5 mm) piece of PDMS containing an 8 mm-long channel (380 μm wide × 585 μm high) was punched with a 5-mm tissue punch to make a cylindrical reservoir. A 0.75-mm tissue punch was used to create a horizontal inlet near the top of the reservoir, to accommodate TT-30 tubing. An outlet was made using a 2.5 mm tissue punch, to accommodate polysiloxane tubing (0.5 mm I.D., 2.2 mm O.D., BioChemFluidics). All tissue punches were from World Precision Instruments, (Sarasota FL, USA). The PDMS layer was plasma bonded to a 1-mm thick glass slide to close the channel, and a flat piece of PDMS was plasma bonded to the open top of the reservoir to close it.

For overnight perfusion of patterned microdevices, a bubble trap was connected upstream of each microdevice after patterning was complete, by inserting TT-30 tubing into the polysiloxane tubing. Chips and bubble traps were placed in a humidified incubator (37 °C, 5% CO_2_) for overnight perfusion. Flow was controlled via an Ismatec IPC-N ISM937C Digital Peristaltic Pump (Cole Palmer, Inc.), using LMT-55 tubing (I.D. 0.25 mm; Cole Palmer, Inc.) and a flow rate of 1.2 μl/min.

### I. Photopatterning and analysis of human lymphocytes (GelSH)

CD4+ T cells were labelled using 10 μM NHS-rhodamine for 20 min at room temperature, rinsed in 1x PBS to remove excess dye by centrifugation at 400 × g for 5 min and resuspended in AIM V serum-free medium (Gibco; Thermo Fisher Scientific, Inc.) supplemented with 10 ng/mL recombinant human IL-7 (R&D Systems; Bio-techne, Inc.) until use.

For micropatterning, cells were resuspended in precursor solution at 1.5 × 10^7^ cells/mL. The 8 arm PEG-NB was added to the precursor to a final concentration of 0.313 mM or 1.25 mM for final concentrations of 2.5 or 10 mM norbornene, respectively, immediately before filling the syringe. Cells were flowed into the device for 2 min at 5 μL/min and photo-patterned as described above. Micropatterned cultures were incubated in a cell culture incubator (37 °C, 5% CO_2_) for 12 hours under continuous perfusion of media (AIM-V, supplemented with 10 ng/mL recombinant human IL-7) at 1.2 μL/min.

After the culture period, the viability of cells was assessed by flowing in a staining solution of Calcein-AM (10 μM) and DAPI (1 μM) in 1x PBS for 2 min at 5 μL/min, which was incubated on-chip for 20 min at 37 °C, then rinsed out for 10 min with 1x PBS at 5 μL/min using a syringe pump. Images were collected using a Zeiss AxioZoom microscope, collecting two to four focal planes per location. Data analysis was performed in ImageJ, as follows: The z-stack images from each location were stacked and converted into a Max Intensity Projection. Cells were identified by using the Particle Analyzer tool (circularity 0.5 – 1, size 12.5 – 500 μm^2^). The percent of live cells was quantified as Calcein-positive cells/ (Calcein-positive + DAPI-positive cells). In preliminary experiments, we confirmed that this concentration of DAPI was low enough to label only the dead cells, and did not double-label Calcein-positive cells.^59,60^

To quantify cell density inside and outside of the patterned structures, images were analyzed using ImageJ. Cells were identified by using the Particle Analyzer tool (circularity 0.5 – 1, size 12.5 – 500 μm^2^). Cell density was calculated as the number of cells per unit area (mm^2^), by selectively analyzing the area inside of all hydrogel features and the negative space in the chamber outside of the features.

### J. Statistical Analysis

Statistical tests and curve fits were performed in GraphPad Prism 8.4.3 and 9.0.2.

## Supplementary Material

The supplementary material file contains supporting figures S1 – S5. S1: Rheological characterization of GelMA and GelSH; S2: Images of optimization of air plasma treatment on PDMS; S3: Schematic of alignment markers on PDMS and photo-mask; S4: Characterization of pattern fidelity under cell culture conditions; S5: Characterization of isolated human naïve CD4+ T lymphocytes.

## Acknowledgments

This work was supported by a seed award from the UVA Center for Advanced Biomanufacturing, and by the National Institute of Biomedical Imaging and Bioengineering (NIBIB) under Award Number U01EB029127 through the National Institutes of Health (NIH), with co-funding from the National Center for Advancing Translational Sciences (NCATS). JOC was supported in part by the NIH-funded T32 Biotechnology Training Program at the University of Virginia. JMZ was supported in part by the Graduate Research Fellowship Program through the National Science Foundation. ANM was supported in part by the Harrison Undergraduate Research Award through the Office of Undergraduate Research (OUR) Grant at the University of Virginia. The authors would like to thank the Lampe lab for early feedback, providing LAP photoinitiator, and access to their lyophilizer, as well as the Landers lab for access to the CO_2_ laser etcher. We would like to also thank the UVA Center for Advanced Biomanufacturing (CAD Bio) for access to the rheometer and the Caliari lab for training and access to the lyophilizer equipment. We thank Amirus Salaheen and Tochukwu Ozulumba for technical assistance during experiments.

## Data Availability

The data that support the findings of this study are available from the corresponding author upon reasonable request.

## Author Contributions

JEOC, JMZ, and RRP led the experimental design, interpreted the results, and drafted the manuscript. JEOC performed, collected and analyzed data for all experiments. JMZ contributed to experimental set-up, data collection, and characterization of biomaterials. AA isolated, characterized, and optimized culture conditions for CD4+ T cells. ANM contributed to fabrication of microfluidic chips in the early stages of the project. JMM provided expertise on analysis of biomaterials. CJL provided expertise for design of experiments with T cells. All authors critically revised the manuscript.

