## Supplemental Information for "Generation of nonlinear and spatially-organized 3D cultures on a microfluidic chip using photoreactive thiol-ene and methacryloyl hydrogels"

Electronic Supplementary Materials for

University of Virginia

Charlottesville, VA 22904, USA

**Contents:**

- Supporting Figure S1: Rheological characterization of GelMA and GelSH
- Supporting Figure S2: Images of optimization of air plasma treatment on PDMS
- Supporting Figure S3: Schematic of alignment markers on PDMS and photo-mask
- Supporting Figure S4: Characterization of pattern fidelity under cell culture conditions
- Supporting Figure S5: Characterization of isolated human naïve CD4+ T lymphocytes

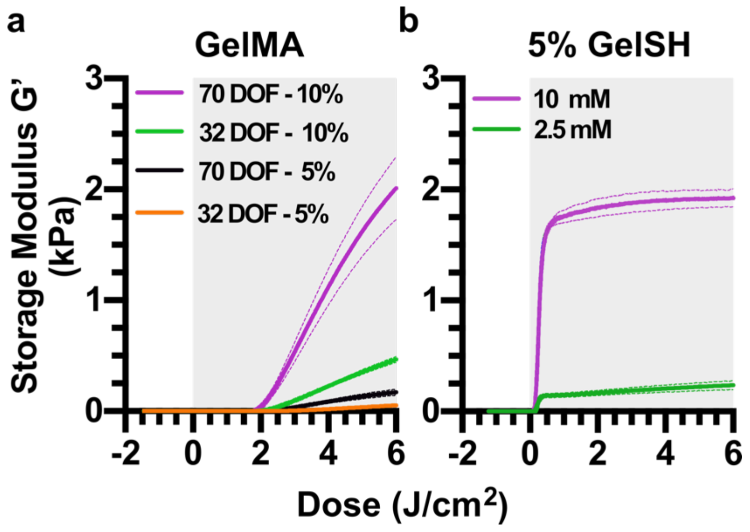


**Fig. S1** Initial testing of hydrogel formulation to access biomimetic storage moduli. (a) Rheometry measurements of the storage modulus of the GelMA during *in situ* polymerization under constant light exposure at 50 mW/cm$^2$. Legend indicates the degree of functionalization (DOF, %) of the gelatin and concentration (% w/v) of GelMA present. n=3. (b) Rheometry measurements of the storage modulus of 5% w/v GelSH during *in situ* polymerization under constant light exposure at 50 mW/cm$^2$. Legend indicates the concentration of norbornene, where there is 8 mol norbornene per mol PEG-NB. Lines show mean (solid) and std deviation (dashed), n=3 technical replicates. Grey shading indicates the time period when the light was turned on.

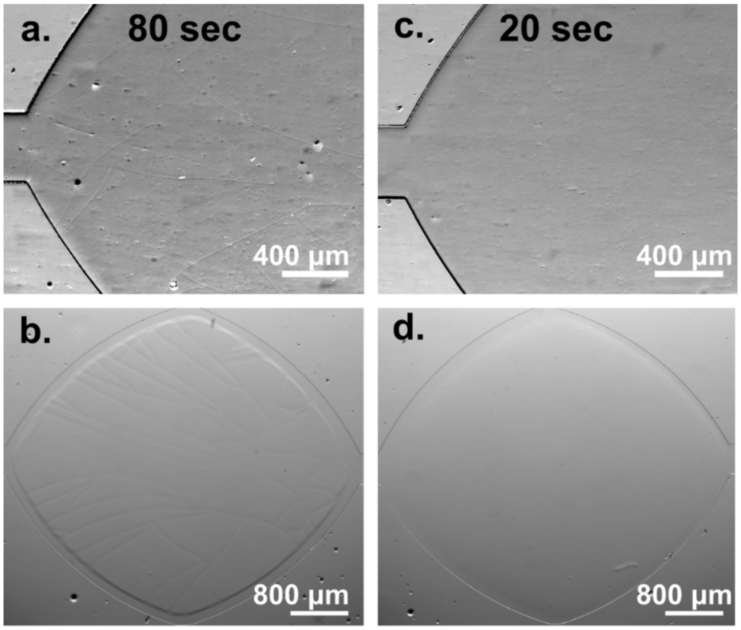


**Fig. S2 Optimization of exposure time to air plasma prior to bonding of the microfluidic device.** (a) Zoomed-in transmitted light image of PDMS showed cracking after 80 sec exposure to air plasma. (b) On-chip patterned hydrogel (10% GelNB, 3.4 mM LAP, 3.75 mM 4-arm PEG-SH) exhibited defects that were templated by the cracks in the PDMS. (c) Zoomed-in image of smooth PDMS surface after 20 sec exposure to air plasma. (d) Patterned hydrogel without cracks and/or defects on a chip that was plasma-treated for 20 seconds.

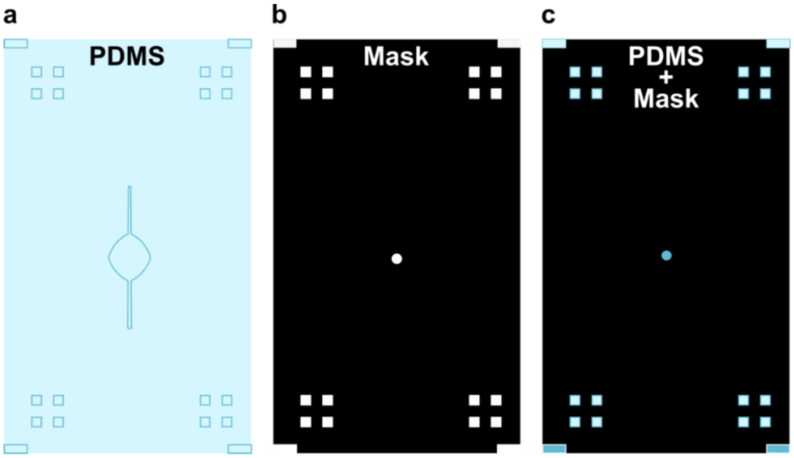


**Fig. S3 Alignment of photomask for photopatterning on chip.** (a) Schematic of the PDMS layer, which included recessed alignment markers in each corner. Alignment marks were transferred to the PDMS from the SU-8/silicon master. (b) Schematic of a photomask with a transparent central circular feature and transparent alignment markers in each corner. (c) Schematic of PDMS layer aligned with photo-mask. Blue tint from PDMS comes through transparent alignment markers in photomask.

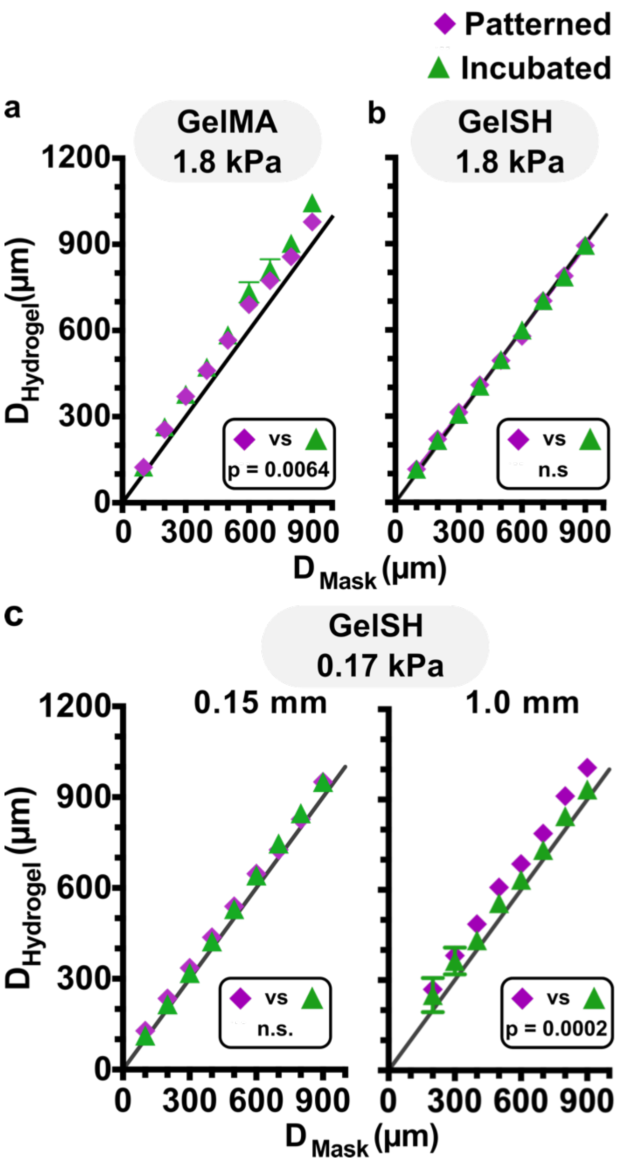


**Fig. S4 Assessing the stability of pattern resolution after incubation.** (a-c) Quantification of accuracy between the diameter of the design on the photomask and the resulting diameter of the hydrogel region, measured either immediately after patterning ("patterned") or after an additional 30-min incubation and rinse ("incubated"). Symbols and error bars represent mean and standard deviation; some error bars too small to see. Significance obtained by Paired T-test, n.s. indicates $p > 0.05$, in all panels. Black line represents $y = x$. (a) data for 1.8 kPa GelMA hydrogel n = 3, p = 0.0064. (b) data for 1.8 kPa GelSH, n = 4, n.s. (e) data for 0.17 kPa GelSH, for microfluidic chips made with (left) a 0.15 mm coverslip, n = 4 chips, or (right) a 1 mm glass slide, n = 3 chips.

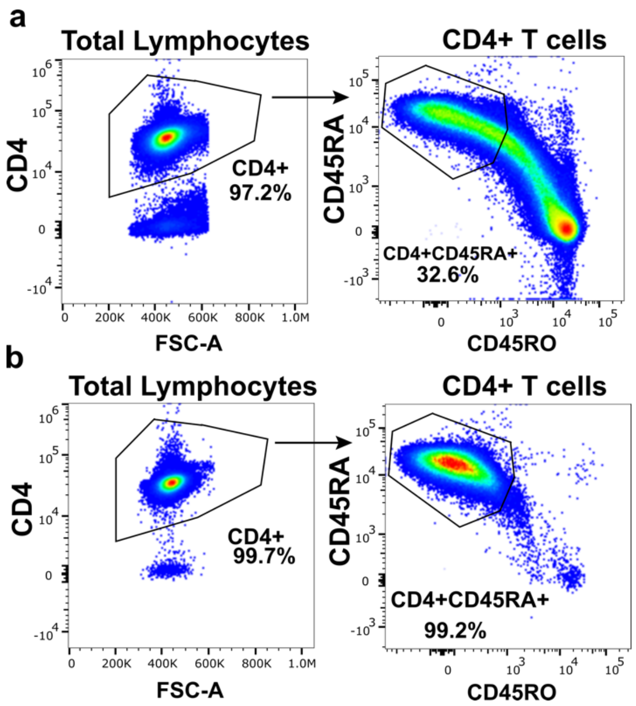


**Fig. S5 Representative characterization of isolated human naïve CD4+ T lymphocytes.** Naïve CD4+ T cells were isolated using a two-step enrichment process from TRIMA collars obtained from healthy donors. Purity of isolated naïve CD4+ T cells was determined using flow cytometry after surface staining of lymphocytes with antibodies against CD4, CD45RA, and CD45RO. (a) Phenotype of the T cell population after RosetteSep™ and Ficoll-Paque density centrifugation, showing enrichment of total CD4+ T lymphocytes. (b) Phenotype of the T cell population after isolation from enriched total CD4+ T cells using EasySep™ negative selection kit, showing high purity of naïve CD4+ T lymphocytes (CD4+CD45RA+CD45RO-)**.**
